# Directional predictions of HIV transmission with optimised genomics in cohorts of serodiscordant couples

**DOI:** 10.1101/2023.10.19.563197

**Authors:** Lele Zhao, Matthew Hall, Chris Wymant, Lucie Abeler-Dörner, Newton Otecko, George MacIntyre-Cockett, Sandra E. Chaudron, Laura Thomson, Tanya Golubchik, Jairam Lingappa, Luca Ferretti, Katrina Lythgoe, Christophe Fraser, Joshua Herbeck, David Bonsall, the PANGEA consortium

## Abstract

Viral genetic information from people living with HIV can deepen our understanding of the infection’s epidemiology at many scales. To better understand the potentials and limits of tools that utilise such information, we show the performance of two representative tools (*HIV-TRACE* and *phyloscanner*) in describing HIV transmission dynamics, with different types of genetic data, and compare with previous findings. The samples were collected from three cohort studies in Sub-Saharan Africa and were deep sequenced to produce both short Illumina reads and long PacBio reads. By comparing *phyloscanner*’s performance with short and long reads, we show that long reads provide improved phylogenetic resolution for the classic transmission topology in joint within-host trees. Our pipeline accurately predicted the direction of transmission 88%-92% of the time. We also show that the timing of sample collection plays an important role in the reconstruction of directionality using deep sequencing data. Consensus sequences were also generated and used as *HIV-TRACE* input to show different patterns of clustering sensitivity and specificity for data from different genomic regions or the entire genome. Finally, we discuss adjusting expectations about sensitivity and specificity of different types of sequence data, considering rapid pathogen evolution, and highlight the potentials of high within-host phylogenetic resolution in HIV. In conclusion, viral genetic data collected and presented differently could greatly influence our ability to describe the underlying dynamics. Methods for source attribution analysis have reached levels of superior accuracy. However, residual uncertainty emphasizes sequence analysis alone cannot conclusively prove linkage at the individual level.

**Importance:** Understanding HIV transmission dynamics is key to designing effective HIV testing and prevention strategies. By using different sequencing techniques on well-characterised cohorts, we were able to evaluate the effect of genetic data resolution on the accuracy of identifying likely transmission pairs and the direction of transmission within pairs. We find that the longer reads generated by PacBio sequencing are more suitable for transmission analyses.

## Introduction

Good progress has been made towards the UNAIDS 2020 90-90-90 testing and treatment target (90% of infected individuals knowing their HIV status, 90% of those on treatment, and 90% of those virally suppressed), even if the 2020 target was missed (UNAIDS 2020). In setting its even more ambitious target of 95-95-95 by 2025, UNAIDS emphasised the need for person-centred and context-specific support services (UNAIDS 2020, 2021). In epidemics in which incidence is declining, it becomes more and more important to understand where transmissions still occur and how those people living with HIV who have not yet been reached by HIV services can be linked to care.

Pathogen transmission analyses using viral genetic data are useful tools for HIV epidemic control. Consensus sequences are widely used in studies of pathogen transmission dynamics. Analysis of consensus-level genomic sequences, where the consensus can be thought of as the representative sequence of a sample, can provide overall population statistics such as the number of clusters and their size (Wertheim et al. 2018; Bezemer et al. 2022), drug resistant mutation prevalence (Blassel et al. 2021; Dalmat et al. 2018), and risk factors associated with sources of transmission (Reichmuth, Chaudron et al. 2021; Hué et al. 2014; Le Vu et al. 2019).

Next-generation sequencing (NGS) data provides detailed sequence information about the within-host viral population while the consensus is a summary of that. NGS data usually consist of large number of short reads produced by platforms such as Illumina or by platforms which generate longer continuous reads, such as PacBio and Oxford Nanopore. They can be used to understand within-host viral dynamics (Zanini et al. 2015; Thys et al. 2015; Raghwani et al. 2018) and more acutely describe transmission dynamics in a population. These various types of NGS data can facilitate analyses to elucidate within-host selection and population structure (Illingworth et al. 2020; Raghwani et al. 2019), to evaluate drug resistance pre-treatment (Dalmat et al. 2018; Dauwe et al. 2016), to study the correlations hitchhiking between drug resistance mutations and linked resistance-associated mutations (Flynn et al. 2015), to provide increased resolution to genomic source attribution (Rose et al. 2019; Zhang et al. 2021; Hall et al. 2021), and to give information about the dynamics of the transmitted/founder lineages (Kijak et al. 2017; Le et al. 2015). Longer continuous reads are more informative for linking neighbouring mutations, which can increase the power of phylogenetic analyses and facilitating haplotype reconstruction (Nguyen Quang et al. 2020; Mori et al. 2022; Laird Smith et al. 2016).

Determining the likely direction of transmission gets to the heart of infectious disease epidemiology by allowing the identification of transmission risk factors, not just acquisition risk factors. This may allow prevention and treatment resources to be more effectively concentrated on populations who are at highest risk of transmitting. Some characteristics may be associated with increased transmission in the general population. Others may be associated with increased transmission from subpopulations with particular characteristics, revealing flows of transmission between population subgroups (Le Vu et al. 2019; Reichmuth, Chaudron et al. 2021; Bbosa et al. 2020; Ratmann et al. 2020). For example, an analysis of transmission direction using NGS sequences found that a geographic area with high HIV prevalence was a sink of transmission rather than (as had previously been assumed) a source (Ratmann et al. 2020). Recent studies of HIV in Zambia and Botswana identified men aged 25-40 as a priority demographic for future prevention programmes (Hall et al. 2021; Magosi et al. 2022).

Previous studies which investigated the performance of HIV transmission inference focused on short read NGS sequencing data. One study found NGS sequencing data to be similarly informative to Sanger-sequenced consensus genomes when analysing *pol*-gene fragments to identify transmission chains (Todesco et al. 2019). They noted inconsistency in these methods and cautioned against using them as the sole evidence in HIV transmission studies. Other studies found high sensitivity for inferring the direction of transmission using NGS data (Rose et al. 2019), that reliable population-level inferences could be made from estimates of the direction of transmissions (Ratmann et al. 2019), and that sequences from the *pol* gene were better for inferring direction than sequences from *gag* or *env* (Zhang et al. 2021). They also suggested that phyloscanner is more accurate if the time difference between the index (HIV positive individual) and seroconverter (paired HIV negative individual who later seroconverted) samples is greater.

In this study, we systematically evaluate approaches for identifying highly likely transmission pairs. We used a dataset of enrolled serodiscordant couples with frequent follow-ups from three different cohort studies/trials, where the 247 heterosexual couples with available viral genetic data were self-reported partners and became seroconcordant during the studies, providing a “gold standard” set of transmission pairs for the comparison of genomic clustering and source attribution methodologies. We provide a detailed demonstration of the impact of the type of sequencing data (consensus, short-reads and long-reads), sampling strategy and two software tools (*HIV-TRACE* and phyloscanner) on the accuracy of linkage and directionality of the likely transmission pairs. HIV-TRACE, a commonly used and highly scalable tool, uses unaligned viral consensus sequences to constructs networks based on pairwise genetic distance with a given threshold (Kosakovsky Pond et al. 2018). The package phyloscanner (Wymant, Hall, et al. 2018) is a software tool for analysing between- and within-host pathogen dynamics using NGS data by constructing alignments and phylogenies within user-defined genomic windows. Unlike HIV-TRACE, phyloscanner can be used to identify probable transmission pairs and their likely direction of transmission. In this study, we found more variability in the sensitivity and specificity of clustering transmission pairs using different regions of the consensus sequences compared to short-read NGS data. We also found that the time of sampling and the length of the NGS reads all impact the phylogenetic resolution for calling the direction of transmission.

## Methods

### Study cohorts

We analysed 247 heterosexual couples from three studies, with longitudinal samples available for 235 of the couples. 16 pairs were from the Cohort Observation Study (COS, Lingappa et al. 2011), which recruited 485 serodiscordant couples from South Africa and Uganda without interventions and followed them up quarterly for 12 months. 120 pairs were participants of the Partners in Prevention PrEP study. This randomised, double-blinded, placebo-controlled trial recruited 4758 serodiscordant couples from Kenya and Uganda, and followed them up for 24-36 months (quarterly for the infected partner, and monthly for the susceptible partner) (Mujugira et al. 2011; Baeten et al. 2012). This study took place from July 2008 to November 2010. The final 111 couples were participants of the Partners in Prevention HSV/HIV study (Lingappa et al. 2009; Celum et al. 2010). This randomised, double-blinded, placebo-controlled trial recruited 3408 couples in East and Southern Africa and followed them up for a maximum of 24 months (monthly for the HIV-1 infected partner, and quarterly for the susceptible partner). This study took place from November 2004 to April 2007. Study participants were not on HIV antiviral treatment at enrollment and during the trials, as per recruitment and trial criteria. All individuals whose samples were analysed in this study provided written informed consent for sample storage and future genetics studies. The University of Washington Human Subjects Review Committee and ethics review committees at local and collaborating institutions of each study sites approved the study protocols (The Partners in Prevention HSV/HIV Transmission Study was registered with ClinicalTrials.gov (#NCT00194519)).

### Sample preparation, sequencing, and data processing

1186 samples were processed using the veSEQ-HIV protocol described in detail previously (Bonsall et al. 2020) and the application of the protocol on the Illumina platform was recently accredited for drug resistance testing in HIV (Jenkins et al. 2023). In brief, total RNA was extracted from HIV-positive plasma. Libraries were prepared using the SMARTer Stranded Total RNA-Seq kit v2 - Pico Input Mammalian (Clontech, TaKaRa Bio). Unfragmented RNA was reverse transcribed with adapter-linked random hexamers and the first strand cDNA was then converted into double-stranded dual-indexed DNA libraries, with a maximum of 12 PCR cycles. Libraries with compatible indexes were pooled and cleaned to eliminate shorter fragments. Custom HIV-specific biotinylated 120-mer oligonucleotide probes (IDT) (Jenkins et al. 2023) were used to capture HIV DNA fragments and the captured libraries were amplified for sequencing on Illumina NovaSeq. 491 of the enriched metagenomic-libraries sequenced on Illumina were repooled and ligated to SMRT-bell Pacbio adapters (SMRTbell® prep kit 3.0, Sequel® II binding kit 2.1) and resequenced on a PacBio Sequel IIe instrument. Size-distributions of individual and pooled libraries were differentially size-fractionated for long and short read sequencing by controlling ratios of PEG and sample volume during DNA cleanups post-PCR, using Ampure-XP beads (0.5/0.68 for Pacbio, 0.68/0.8 for Illumina).

The demultiplexed Illumina sequencing reads were processed with Kraken (Wood and Salzberg 2014) to remove human and bacterial reads. The resulting viral and unclassified reads had adapters and low-quality bases removed using Trimmomatic (Bolger, Lohse, and Usadel 2014). Contigs were assembled with SPAdes (Bankevich et al. 2012) and metaSPAdes (Nurk et al. 2017) with default parameters. Contiguous sequences were clustered using cd-hit-est (Fu et al. 2012) to remove redundant contigs. The processed reads were mapped to sample-specific references constructed from the contigs, and then a consensus sequence was called from the mapped reads, using *shiver* (Wymant, et al. 2018). For samples for which no contigs could be assembled, the reads were compared to a set of 199 HIV reference genomes from the Los Alamos HIV database (http://www.hiv.lanl.gov) by Kallisto (Bray et al. 2016) to find the closest matching genome and were subsequently mapped to this reference genome using *shiver*. Deduplication of mapped reads using Picard (https://broadinstitute.github.io/picard/) was enabled in *shiver*. The same bioinformatic pipeline developed for paired-end short-read data was adapted for longer-read Pacbio high-fidelity (HiFi) reads (closed circular-consensus sequences; ccs). These reads were demultiplexed using a bespoke configuration of the Pacbio LIMA tool and a dummy reverse complement-read was synthesised to mimic paired-read data (required by *shiver*).

### Dataset organisation by time since infection

#### 1. Determination of time of transmission/infection

We estimated the time between infection and collection for each sample using HIV-phyloTSI (Golubchik et al. 2022). An estimated time of transmission for each sample was calculated by subtracting the HIV-phyloTSI point estimate of time since infection from the date that sample was obtained. These were combined with the known epidemiological information (the last negative date and first positive date of the seroconverter) to estimate one time point of transmission for each index-seroconverter (i.e. source-recipient) pair (for details please see Supplementary TextFile 1).

#### 2. Generating index-seroconverter sample groups

As illustrated in Figure 1, index samples were classified into three groups by sampling day: pre-transmission (more than 120 days before transmission), around-transmission (within 120 days before or after transmission), and post-transmission (more than 120 days after transmission). Similarly, seroconverter samples were classified into two groups: early (within 270 days after transmission) and late (more than 270 days after transmission). We also included a “multiple” category in which all available samples from the relevant individual were used in the analysis. The time point separating the index and seroconverter sample groups were selected with reference to the sampling time distributions (Supplementary Figure 1-2), to maximise estimation accuracy. Sampling time pairings within a couple are one-to-one, one-to-multiple, or multiple-to-multiple (i.e. first two columns of Table 1) depending on availability.

**Figure 1.**
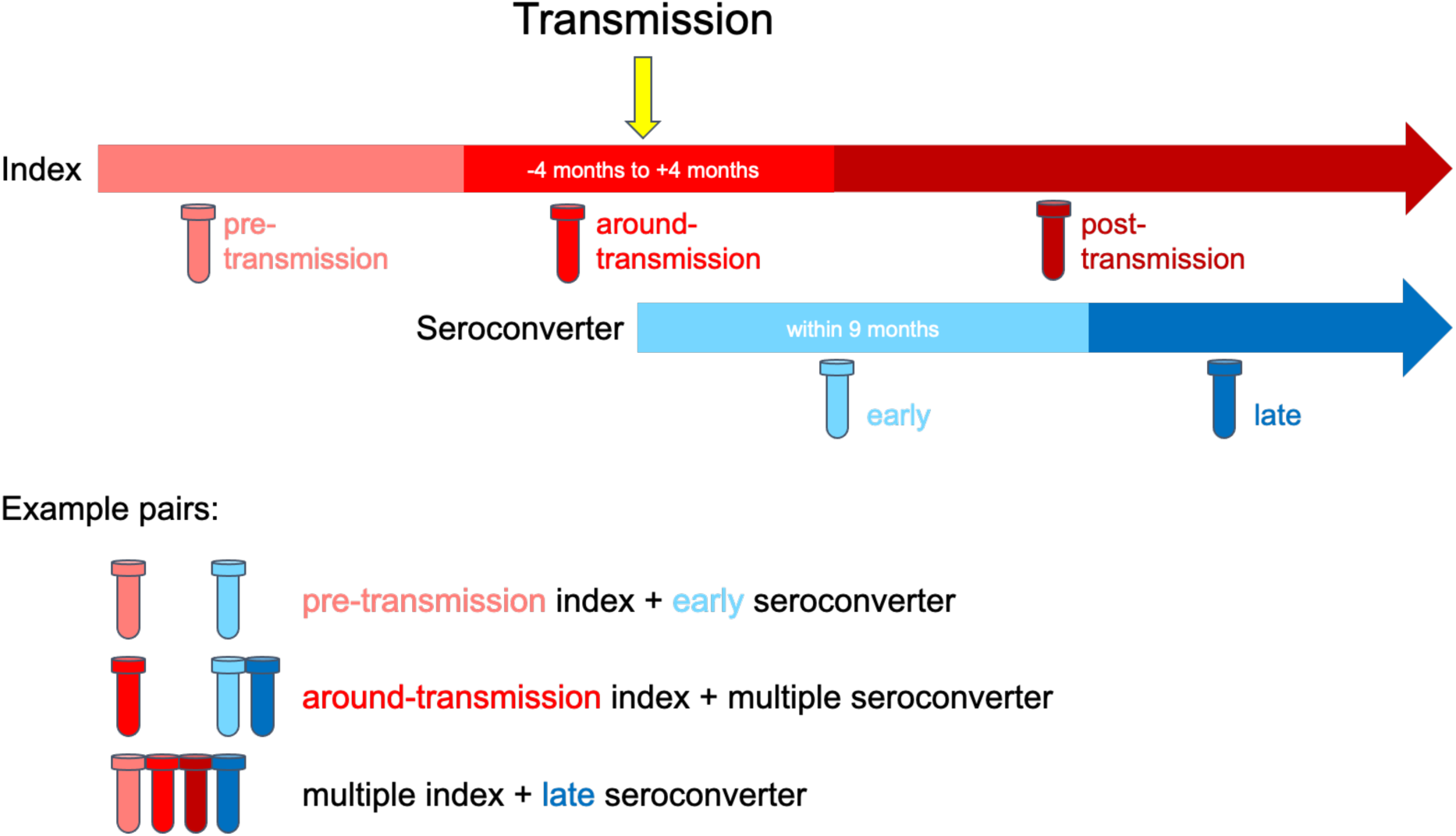
Schematic for classification of index-seroconverter sample pairs.

**Table 1.**
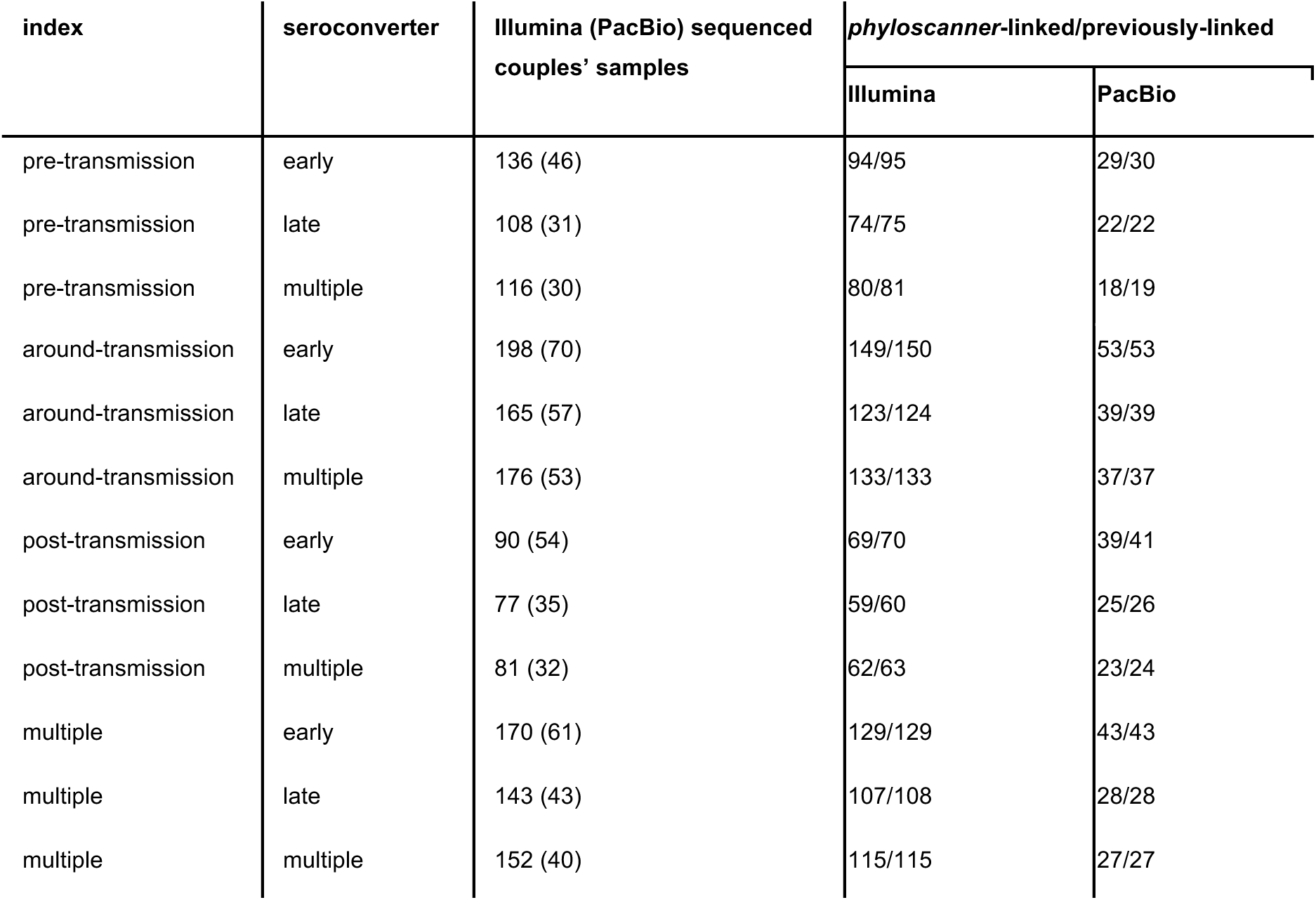
Comparison of the number of linked pairs within index-seroconverter sampling time pairings.

| index | seroconverter | Illumina (PacBio) sequenced couples' samples | phyloscanner-linked/previously-linked |  |
| --- | --- | --- | --- | --- |
|  |  |  | Illumina | PacBio |
| pre-transmission | early | 136 (46) | 94/95 | 29/30 |
| pre-transmission | late | 108 (31) | 74/75 | 22/22 |
| pre-transmission | multiple | 116 (30) | 80/81 | 18/19 |
| around-transmission | early | 198 (70) | 149/150 | 53/53 |
| around-transmission | late | 165 (57) | 123/124 | 39/39 |
| around-transmission | multiple | 176 (53) | 133/133 | 37/37 |
| post-transmission | early | 90 (54) | 69/70 | 39/41 |
| post-transmission | late | 77 (35) | 59/60 | 25/26 |
| post-transmission | multiple | 81 (32) | 62/63 | 23/24 |
| multiple | early | 170 (61) | 129/129 | 43/43 |
| multiple | late | 143 (43) | 107/108 | 28/28 |
| multiple | multiple | 152 (40) | 115/115 | 27/27 |

### Sequence clustering by HIV-TRACE

Consensus genome sequences from shiver, with coverage of at least 7500bp of the HIV-1 genome, were aligned to the HXB2 reference genome with MAFFT v7.490 (Katoh and Standley 2013) using the FFT-NS-2 (default) strategy, and cut into *gag*, *pol* and *env* gene blocks using the HXB2 coordinates (*gag*: 790-2292, *pol*: 2358-5096, *env*: 6225-8795). Sequences of *gag*, *pol*, *env* and the whole genome served as inputs for *HIV-TRACE* (additional options used available in Supplementary TextFile 1), ran multiple times with distance thresholds ranging from to 0.07 substitutions per site with a 0.005 increment. Because the consensus dataset consists of a mixture of subtypes A1, C, D and other genomes, we also supplied subtype A1, C and D sequences as the reference genome in parallel *HIV-TRACE* runs. Different index-seroconverter sampling time pairings’ *HIV-TRACE* runs used sequences contributed by the corresponding samples.

### Gold standard classifier for sensitivity and specificity

To examine the sensitivity and specificity of *HIV-TRACE* in clustering the heterosexual couples on different sampling time pairings, distance thresholds and genomic regions, we relied on a “gold standard classifier”, a linkage assessment method from previous work (Campbell et al. 2011), that identifies “gold standard transmission pairs”. Gold standard transmission pairs are genetically linked couples in this study. All heterosexual couples in the current study determined to be in or not in the “gold standard transmission pairs” set make up our positive or negative set in the sensitivity and specificity calculations. A pair was considered linked by *HIV-TRACE* when at least one sample from each individual was present within the same cluster.

### Linkage and directionality confirmation by phyloscanner

All processed samples with consensus genomes covering at least 7500 base pairs (bp) of the HIV-1 genome were analysed using *phyloscanner*. Each analysis involved one sampling time pairing of one heterosexual couple (for example pairs, see Figure 1). First, we used *phyloscanner* to extract all mapped reads that fully span a given genomic window, and to align all unique sequences among those reads, in overlapping sliding windows along the genome, as previously described (Wymant, Hall, et al. 2018) (we defined window coordinates with respect to the HXB2 reference sequence; the exact phyloscanner commands are given in Supplementary TextFile1). For samples sequenced with the Illumina platform we used windows of width 251bp starting every 10bp (i.e. overlapping by 240bp), while for those sequenced with the PacBio platform the windows were of width 1501bp starting every 60bp (i.e. overlapping by 1440bp). We also examined reads in the *gag*, *pol* and *env* regions separately; for these, only windows that fall entirely within the gene’s HXB2 coordinates (specified earlier) were included. The phylogeny-building step of *phyloscanner* used IQ-TREE 1.6.12 (Nguyen et al. 2015) with the substitution model GTR+F+R6. Known drug resistance sites (listed in Supplement TextFile 1) in HXB2 were masked to prevent the tree topology from being affected by potential homoplasy at shared drug resistance sites.

For the second step of *phyloscanner* (command in Supplement TextFile1), which reconstructs transmission, linkage of two individuals was considered confirmed when at least half of all available genomic windows showed normalised minimal patristic distance between reads from the two individuals to be equal or less than a distance threshold. The direction of transmission was considered confirmed when the fraction of windows showing phylogenetic support for index→seroconverter was equal to or above a set threshold, and the fraction of seroconverter→index windows was below the same threshold. Table 2 outlines the criteria for the four inferred directions of transmission at a threshold of 33.3%. *Phyloscanner*’s sensitivity and specificity in linking transmission pairs were also examined.

**Table 2.** Criteria for inferring transmission direction using *phyloscanner*.

| Transmission direction of a pair | index→seroconverter | seroconverter→index | conflicting | unconfirmed |
| --- | --- | --- | --- | --- |
| Fraction of windows index→seroconverter | ≥33.3% | <33.3% | ≥33.3% | <33.3% |
| Fraction of windows seroconverter→index | <33.3% | ≥33.3% | ≥33.3% | <33.3% |

## Results

### Linkage of transmission pairs from consensus and NGS sequences

We used the consensus and NGS sequences of 1186 samples from 247 heterosexual couples to determine likely transmission. Whole-genome consensus sequences, *gag*, *pol* and *env* alignments were used as input to *HIV-TRACE*. Illumina short reads were used in linkage sensitivity and specificity analyses with *phyloscanner*. The samples were divided into the previously described sampling time pairings for additional testing. In short, *phyloscanner* showed stable and high sensitivity and specificity in linking transmission pairs across distance thresholds (Supplementary Figure S3). An expected downward trend of specificity in all sampling time pairings was observed as genetic distance increased. The results from *HIV-TRACE* were more variable. For both index and seroconverter, datasets including multiple samples had higher sensitivity when compared to those comprising only one sample per individual (Supplementary Figure S4). Using *gag* and *pol* showed similar trends in sensitivity and specificity as a function of the genetic distance threshold. Sensitivity for *gag* and *pol* plateaued between 0.02-0.04, with little increase as the threshold increased, and specificity dropped at shorter genetic distances for the *pol* gene than for the *gag* gene (Figure 2 & Supplementary Figure S4). This could be because *pol* is more conserved. The sensitivity of *env* sequences in clustering transmission pair sequences showed a relatively steady rise starting from small values at low thresholds. The high specificity when using *env* sequences indicated the divergent nature of the envelope gene and the high within-transmission-pair specificity of this region. Analyses using whole genome consensus sequences showed an averaged effect of the three largest regions of the genome. Although the slope of sensitivity was steeper compared to *gag* and *pol*, the sensitivity almost always plateaued before 0.04 in *HIV-TRACE* distance threshold with whole genome as inputs across the various sampling time pairings. The specificity was maintained above 93% for all distance thresholds below 0.07. Figure 2 shows the results from the analyses using only around-transmission index and early seroconverter samples with A1 as the reference subtype, results are similar for when reference genomes are subtype C and D. The pairwise patristic distance distributions of gold standard classifier determined linked and unliked pairs are also available for reference (Supplementary Figure S5).

**Figure 2.**
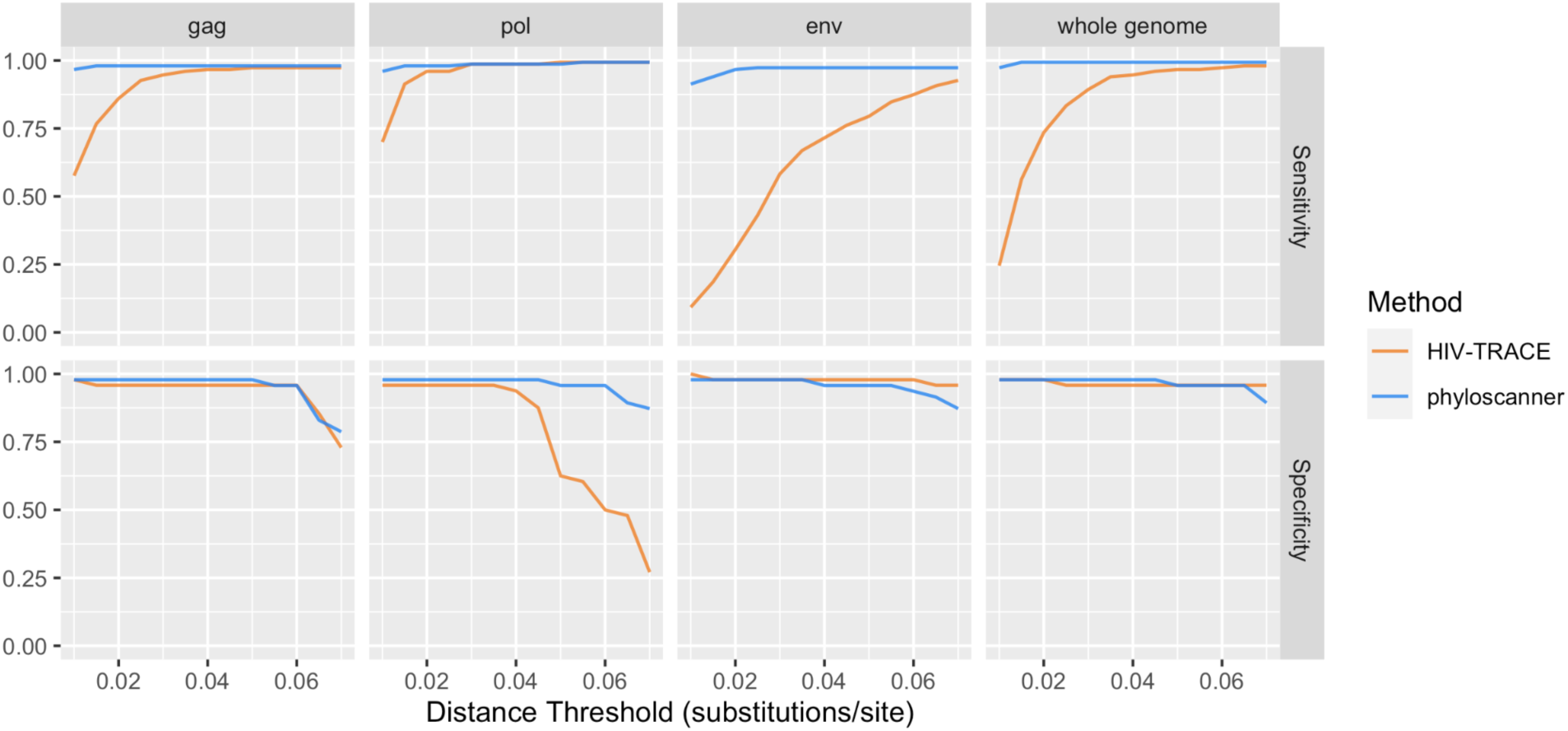
Consensus (HIV-TRACE) and NGS (phyloscanner with Illumina only) sequence data sensitivity and specificity in correctly linking transmission pairs using only around-transmission index and early seroconverter samples. The analysis was performed with gag, pol, env region and whole genome, across genetic distance thresholds 0.01 to 0.07 substitutions/site.

### Confirmation of linkage of transmission pairs with phyloscanner

As PacBio reads were only available for a subset of samples, we summarised the confirmation of transmission linkage using only Illumina sequences with *phyloscanner (distance threshold: 0.02 substitutions/site)*. The mean number of 251bp windows per *phyloscanner* run with Illumina reads was 693, with median 788 [range: 66-887]. The mean number of 1501bp windows per *phyloscanner* run with PacBio reads was 123, median 122 [range:4-129]. When 245 of the 247 heterosexual couples were previously assessed with the “gold standard classifier” for their genetic linkage (Campbell et al. 2011), 184 of them were genetically linked and 61 of them were not. Using all Illumina deep sequencing reads and *phyloscanner*, we found 186 of these 247 pairs were phylogenetically linked and 61 not. Of the 186 *phyloscanner* linked pairs, 183 were also linked in the gold standard, 2 were not, and 1 was not assessed previously. One of the two previously unlinked pairs was linked by *phyloscanner* analysis with 100% of the windows with the normalised minimal patristic distance of the reads from the two individuals shorter than 0.02 substitution per site. The other previously not linked pair involved a potentially problematic seroconverter sample: when only this sample was paired with its corresponding index sample(s), the normalised minimal patristic distance between the reads from genomic windows of the two individuals was above 0.02, however when the problematic sample was not included or was not the only sample from the seroconverter, phylogenetic support for linkage was always observed in more than 74% of the windows. Out of the 61 pairs not linked by *phyloscanner*, 59 were also not linked in the gold standard, 1 was linked and 1 was not assessed previously. One pair linked with the previous “gold standard classifier” was unlinked by *phyloscanner* with no window over the 0.02 distance threshold. Collectively, we did not see a systematic difference in linkage confirmation with different sampling time pairings or data types (Table 1). Including multiple samples may dampen effects from the presence of single problematic samples, but not always.

### Calling of direction of transmission pairs with phyloscanner

Within each sampling time pairing, the pairs that were genetically linked by *phyloscanner* were further assessed for their within-host phylogenetic signal of transmission direction from index to seroconverter (Figure 3, Supplementary Figure S6). The correct transmission direction (index→seroconverter) was confirmed using the windows fraction threshold: 33.3%, specified in Methods. In general, using PacBio long-read sequences resulted in higher rates of correct calls and lower rates of incorrect calls in the direction of transmission than using Illumina short-read sequences. Comparing between the seroconverter samples, the direction of transmission was most often correctly called in pairings containing seroconverter late samples, regardless of pairing with single or multiple index samples or read length. Where index sample(s) were concerned, sampling time pairings with pre-transmission index samples and multiple index samples showed higher proportions of index→seroconverter directed transmission pairs. This suggests that earlier samples for the index and later samples for the seroconverter was the optimal sample combination to confirm transmission direction using phylogenetic topology. We also show that using multiple samples from the seroconverter may result in lower success rates of correct calls and higher rates of incorrect calls, especially when working with shorter reads from the Illumina platform; longer reads could be a partial solution to this problem. Using multiple samples from the seroconverter did not provide an advantage when compared to using single samples, as might have been otherwise expected. This was not the case for multiple index samples. For pairings including single index samples with short reads, we saw an increased proportion of the incorrect (seroconverter→index) direction being supported compared to including multiple index samples in the analyses.

**Figure 3.**
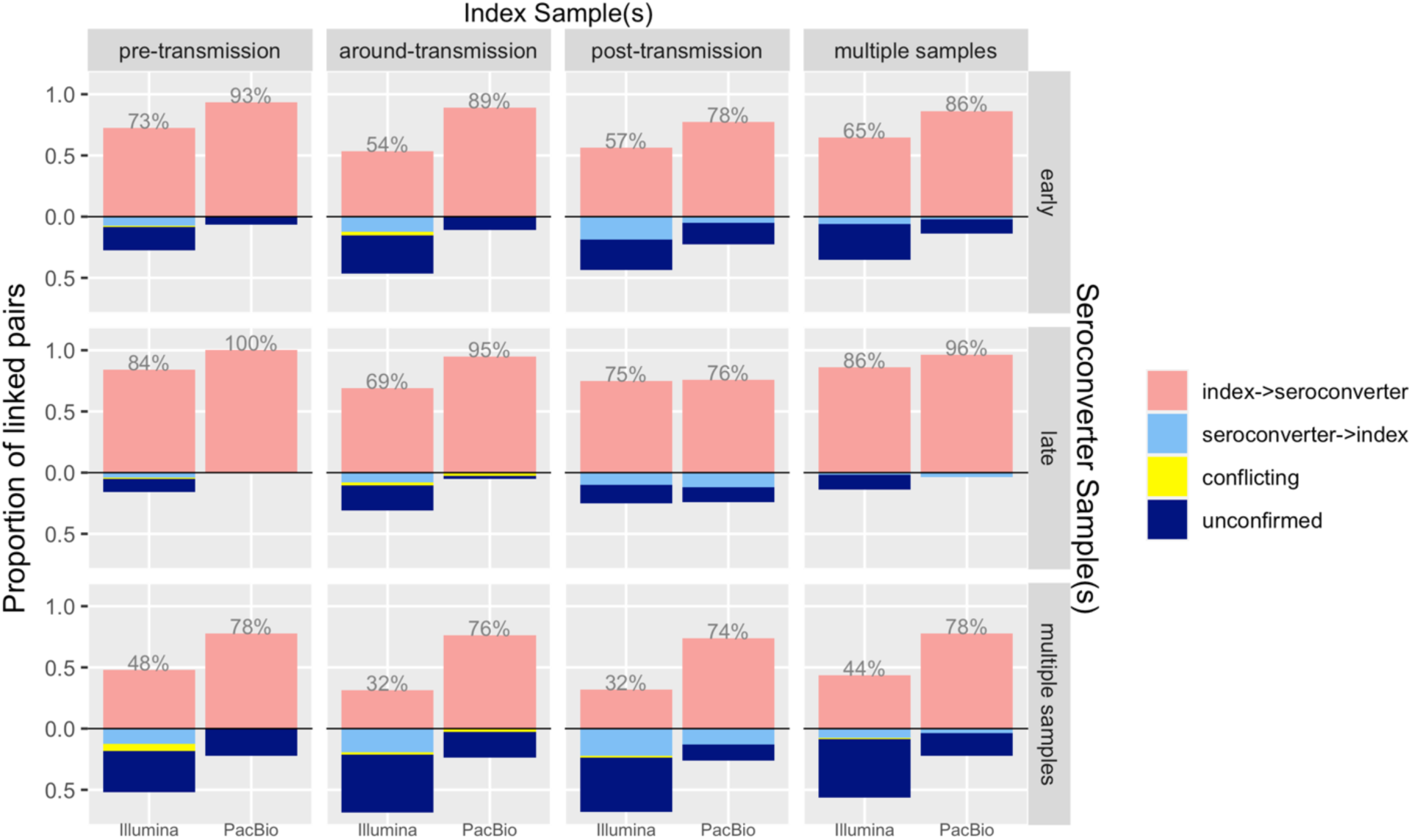
Confirmation of transmission direction by phyloscanner with whole-genome NGS dataset. The columns are index sampling time groups (pre-transmission, around-transmission, post-transmission, and multiple samples) and the rows are seroconverter sampling time groups (early, late, and multiple samples). Within each box, the first column (Illumina) shows results from all available Illumina sequenced pairs, the second column (PacBio) shows results from all PacBio sequenced pairs. The proportion of index→seroconverter (correct calls) are in pink and shown as number labels, seroconverter→index (incorrect calls) are in light blue, conflicting calls are in yellow, and unconfirmed calls are in dark blue.

Using Illumina data from different genomic regions to call transmission direction showed a variable pattern (Supplementary Figure S7). In general, the using sequences from the *env* gene led to higher rates of correctly calling the direction in all sampling time pairings, except when post-transmission index sample and late seroconverter samples were used. The *pol* gene had slightly lower rates of correctly calling direction across all sampling time pairings.

We also varied the fraction of windows used as the threshold for inferring the direction of transmission (Figure 4a, Supplementary Figure S8) to show the effect of this on direction calling using Illumina and PacBio reads. With Illumina reads, the proportion of linked pairs called correctly increased from low thresholds and started to decrease around 0.25 as the threshold continued to increase. For PacBio reads, the proportion of correct calls remained stably high with a slower decline as threshold increases. Long reads enabled high rates of transmission direction calling by increasing the phylogenetic resolution of the joint within-host trees (Figure 4b).

**Figure 4a.**
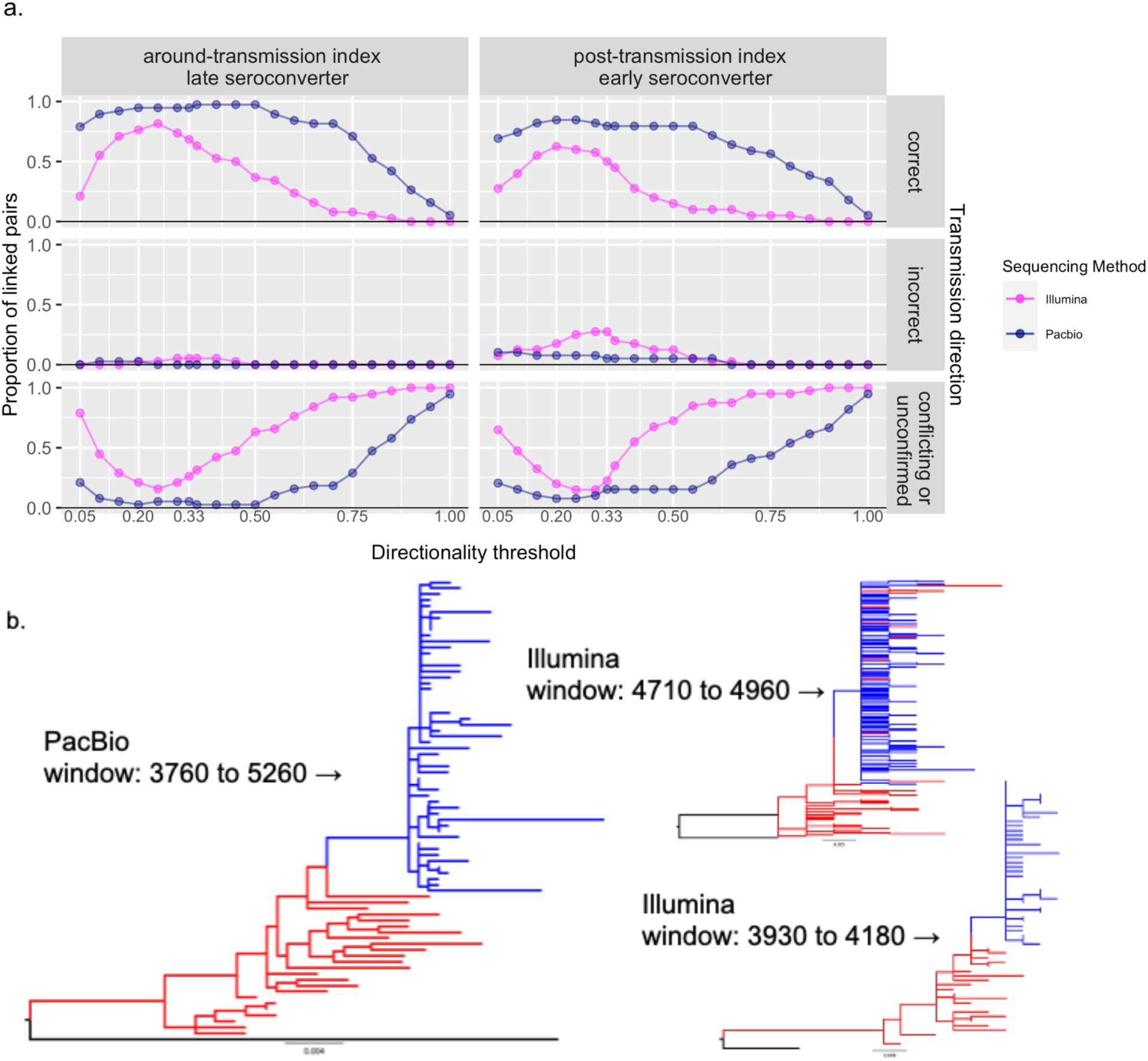
The proportion of phyloscanner linked paired with transmission direction called correctly, incorrectly, and conflicting or unconfirmed across different thresholds. Results from Illumina reads are colored pink, results from PacBio reads are colored dark blue. 4b. Example joint within-host trees built with long PacBio reads from a 1501bp window (HXB2 position 3760 to 5260) and short Illumina reads from 251bp windows (HXB2 position 4710 to 4960 and 3930 to 4180). The branches leading to root/reference sequence (subtype A1) are colored black, index reads are colored red, seroconverter reads are colored blue.

In a further step, we summarised the tree topologies of each directed (Supplementary Figure S9a) and undirected (Supplementary Figure S9b) pair within each sampling time pairing with PacBio data. While the vast majority of windows for direction-confirmed transmission pairs showed index→seroconverter topology, the picture was very different for the undirected pairs. Most windows showed a “complex” tree topology, meaning there was no clear ancestral relationship between the intermingled sequences from the paired individuals. We observed an increased prevalence of “noAncestry” topology in pairings containing post-transmission index samples, where the clade of the index and the clade of the seroconverter were essentially sister clades (Romero-Severson, Bulla, and Leitner 2016). This is expected because the most-recent common ancestor (MRCA) of the index sequences might not predate the MRCA of the seroconverter sequences in these groups. We also checked if a specific region of the genome was associated with non-index→seroconverter tree topologies, but found no obvious region of this property (Supplementary figure S10).

Going back to the original studies, we summarised the percentage of linked individuals and the proportion of those linked whose direction was also confirmed (Table 3). The overall linkage for the couples in the three studies was confirmed when the 0.02 substitutions/site threshold was passed for at least one sampling time pairing. Similarly, the direction of transmission was confirmed when the 33.3% window fraction threshold was passed for at least one sampling time pairing. The proportion linked closely resembled previous findings where 71.5% of the couples were linked (Campbell et al. 2011). Using Illumina generated sequencing reads and phyloscanner, we were able to also confirm the transmission direction of index→seroconverter for 83-88% of the genetically linked pairs. We were also able to confirm the expected transmission direction for more than 89% of the genetically linked transmission pairs using PacBio sequencing reads when available.

**Table 3.**
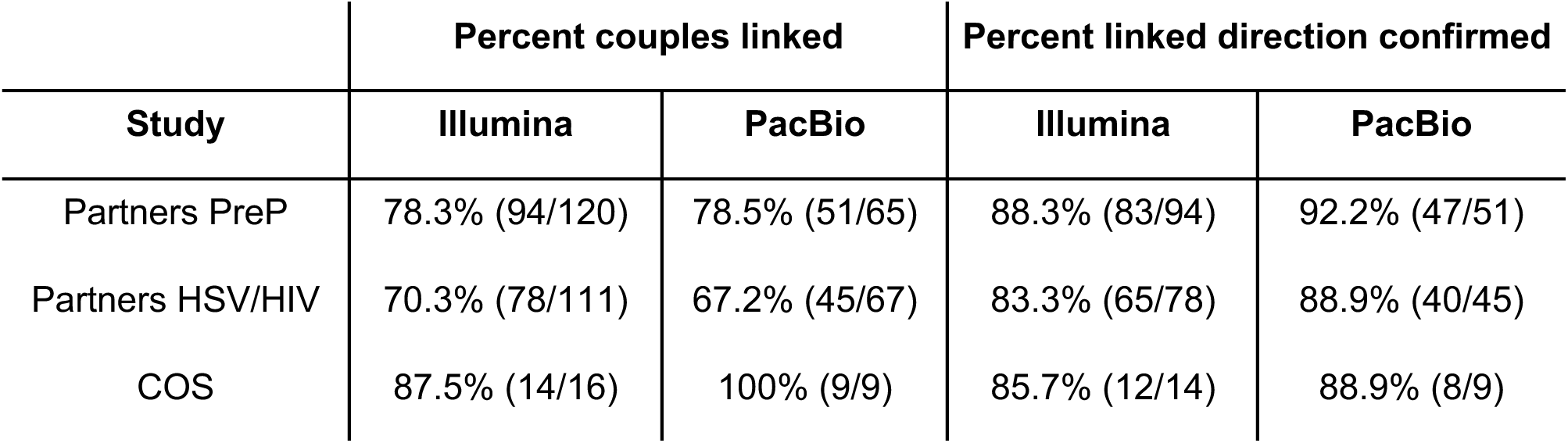
Percentage of genetically linked and direction confirmed heterosexual couples from the three studies.

## Discussion

In this study, we evaluated the effects of using genetic data at different resolutions on investigating linkage (are two individuals a transmission pair) and directionality (which one transmitted to the other) using applicable transmission inference tools in HIV research. With NGS data available for multiply sampled heterosexual serodiscordant couples, we were able to perform sensitivity and specificity analyses to inform future studies of transmission using genetic data. NGS data performed better than consensus-level genetic data which showed variable performance in linking pairs with different HIV genetic regions. We showed high agreement of *phyloscanner* and previous methods in linking transmission pairs. We found no difference in linking pairs with NGS data using either 251bp or 1501bp read windows (for short and long reads respectively), but longer reads improved the phylogenetic resolution in calling the direction of transmission. We also showed the timing of sample collection plays an important role for calling transmission direction using NGS data. While accuracy in calling transmission pairs is improving, the study also shows that the analysis depends on several factors. High accuracy can be obtained on population level (Ratmann et al. 2020; Hall et al. 2021; Bbosa et al. 2020), but studies need to be designed and conducted with care to ensure individuals and groups are not harmed, criminalised or stigmatised and their privacy is protected at all times (Jamrozik et al. 2023; Mutenherwa et al. 2019; Coltart et al. 2018). On the individual level, sequence data alone can never be sufficient to demonstrate transmission in the absence of 100% sampling.

Together with previous studies (Rose et al. 2019; Ratmann et al. 2019), we have shown that improvements in sequencing methods increase our ability to correctly call the direction in HIV transmission pairs. With lower depth and short coverage of the HIV genome, previous studies correctly predicted the transmission direction 55%-74% (Rose et al. 2019) and 76-78% (Ratmann et al. 2019) of the time, while we used phyloscanner with high depth whole-genome NGS short-read data and predicted correctly up to 88.3% of the time, and reaching as high as 92.2% with longer read.

Our findings partially agree with Zhang et al.’s results that higher accuracy in calling directions is obtained where the time between sampling the index and the seroconverter is longer (Zhang et al. 2021). With a larger longitudinal dataset, we were able to confirm that the greatest time difference produced by combining pre-transmission index samples with late seroconverter samples had the highest rate of success in calling transmission direction for both short and long NGS reads. However, we observed different rates of correctly called transmission direction using short reads from different genomic regions and the whole genome. We found *env* to almost always outperform *gag*, *pol* and whole genome in correct calling transmission direction, whereas they found whole genome and *pol* to outperform *env* and *gag*. One possible reason for this could be the difference in window size. Zhang et al. used 340bp windows and we used 251bp (constrained by the read length used and fragment size distribution resulting from sequencing), and we have shown larger windows improve the ability to call transmission direction. We also masked drug resistance sites in this study, many of which are in *pol*, and this reduced the number of available sites in 251bp genomic windows across *pol*. This could result in reads from *pol* being outperformed by reads from *env* in direction calling. The lower transmission direction confirmation rate with Illumina reads than with PacBio reads can be attributed to the phylogenetic resolution limited by the length of the reads. The advantages of long alignments in correctly inferring the transmission direction was also shown in another modelling study (Villabona-Arenas et al. 2022) and discussed previously <u>(Ratmann et al. 2019)</u>. Here, we show that if we apply the knowledge of time of transmission, estimated from epidemiological or deep sequenced genetic data, we can adjust our expectations and filter for samples that will yield less ambiguous results.

Intuitively, one would expect the ability to call transmission direction to increase with the number of samples used in the analyses. However, the answer is more complicated. The length of the alignment used to reconstruct the phylogenetic relationship has to be taken into consideration for a pathogen as fast evolving as HIV. Phylogenetic uncertainty caused by within-host evolution away from the transmitted/founder variant, as previously suggested (Günthard and Kouyos 2019; Rose et al. 2019). When using shorter reads, we see sampling time pairings with late seroconverter samples performing better in confirming transmission direction than early seroconverter samples. This is to be expected as the later samples are more divergent, which would result in them more likely to cluster monophyletically with a long branch connecting to the index clades. However, it must be noted that with more divergence, the nested clade (paraphyletic-monophyletic) topology suggesting transmission tends to turn into monophyletic-monophyletic topologies and directionality is lost (Romero-Severson, Bulla, and Leitner 2016), as shown in Supplementary Figure S9b in pairings involving post-transmission index samples. While the earlier seroconverter samples are more similar to the index sequences, and with the limitation in sequence length, the expected ancestral relationship of source to recipient may be underpowered. We are not advocating the use of single samples over multiple samples, but would like to flag the potential evolutionary information carried by multiple samples in a phylogenetic reconstruction and the complexity it may bring to result interpretation. A balance between the suitability of the method and the data needs to be considered. In addition, unless closely followed-up by study trials, most population sampling-identified source individual samples are from the post transmission period, because the recipient individuals need also be present to find the source. Therefore, we should expect intrinsic biological limitations to call transmission direction even with detailed NGS data and the available methods.

While we are relying on mono/polyphyletic nesting of seroconverter sequences within the “ancestral” index clades, complex relationship tree topologies still dominated 10% of the linked transmission pairs of the studied cohorts. Although the pairs are genetically linked and are with known partnership by study follow-ups, their within-host phylogenies display much more complex relationships. We have shown that the complex topologies are not completely attributable to sampling time or low phylogenetics resolution. Therefore, further studies are required to characterise the evolutionary processes of these atypical joint within-host trees.

In a study involving genetic distance, such as this one, results for sensitivity are more generalisable than those for specificity. This is because the genomic distances between true positives (transmission pairs) are much less variable than those of true negatives, which are simply pairs of individuals who are not linked by transmission. These can range from people who are just two links removed in the transmission chain to those infected with different subtypes. Any reasonable choice of genetic distance threshold will be considerably more effective at ruling out transmission between pairs of the latter sort than the former. As a result, the calculated specificity of any given threshold cannot be entirely separated from the sampling frame of the study.

Our comparison of the rate of correctly calling the direction of transmission using the Illumina and PacBio platforms is presented as a proof of concept. The sample libraries used on the PacBio platform were not specifically optimised as they were residuals from Illumina runs. Future optimisations to the laboratory protocol may produce more and longer fragments to power HIV phylogenetic studies. Improved ability in calling transmission direction is only one of the many analyses that will benefit from increased phylogenetic resolution of long reads.

In summary, we have shown that different tools, different genomic regions and different sampling strategies may affect the limits to which we can infer transmission. Phylogenetics combined with NGS data has proven to be a powerful approach to studying pathogen transmission dynamics, and clearer interpretations of future results will be made possible by understanding the limits of the data.

## Data Sharing

Consensus sequences will be submitted to GenBank. For further data, please contact the PANGEA consortium (www.pangea-hiv.org).

## Acknowledgments

We thank all the study participants and the PANGEA consortium for making this work possible. This work was supported by a Li Ka Shing Foundation Grant awarded to C.F. Sequencing was performed by the PANGEA consortium and the AMPHEUS study which are funded by the Bill & Melinda Gates Foundation (Grants INV-007573 & INV-003680). The research was supported by the Wellcome Trust Core Award Grant Number 203141/Z/16/Z with funding from the NIHR Oxford BRC. The views expressed are those of the author(s) and not necessarily those of the NHS, the NIHR or the Department of Health.

## Supplementary Figures

**Supplementary Figure S1-S2.**
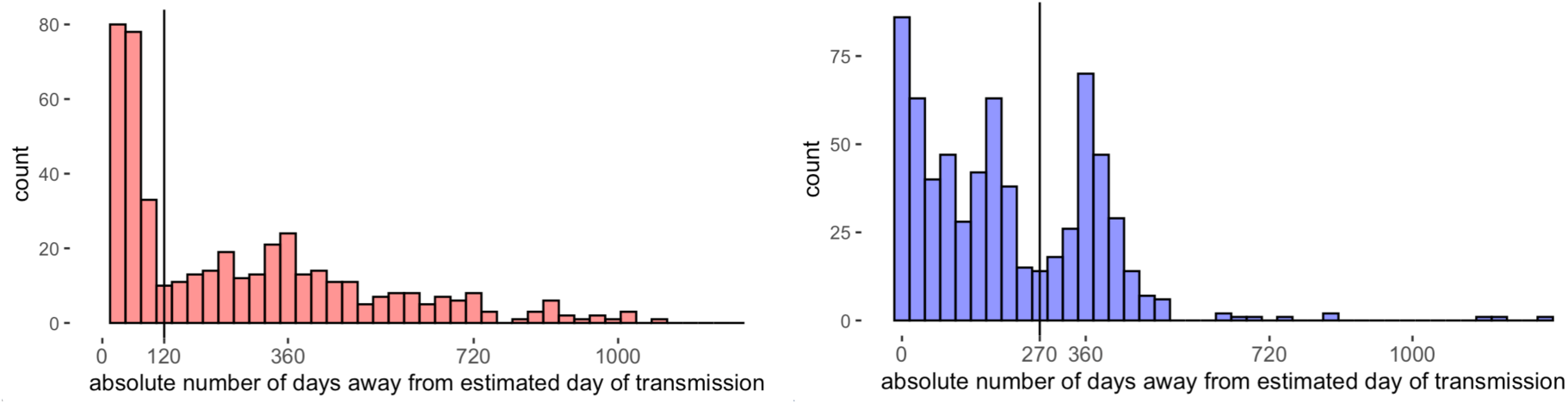
The distribution of sampling times relative to estimated time of transmission. For index samples (red), the vertical line indicates 4 months since transmission. For seroconverter samples (blue), the vertical line indicates 9 months since transmission.

**Supplementary Figure S3.**
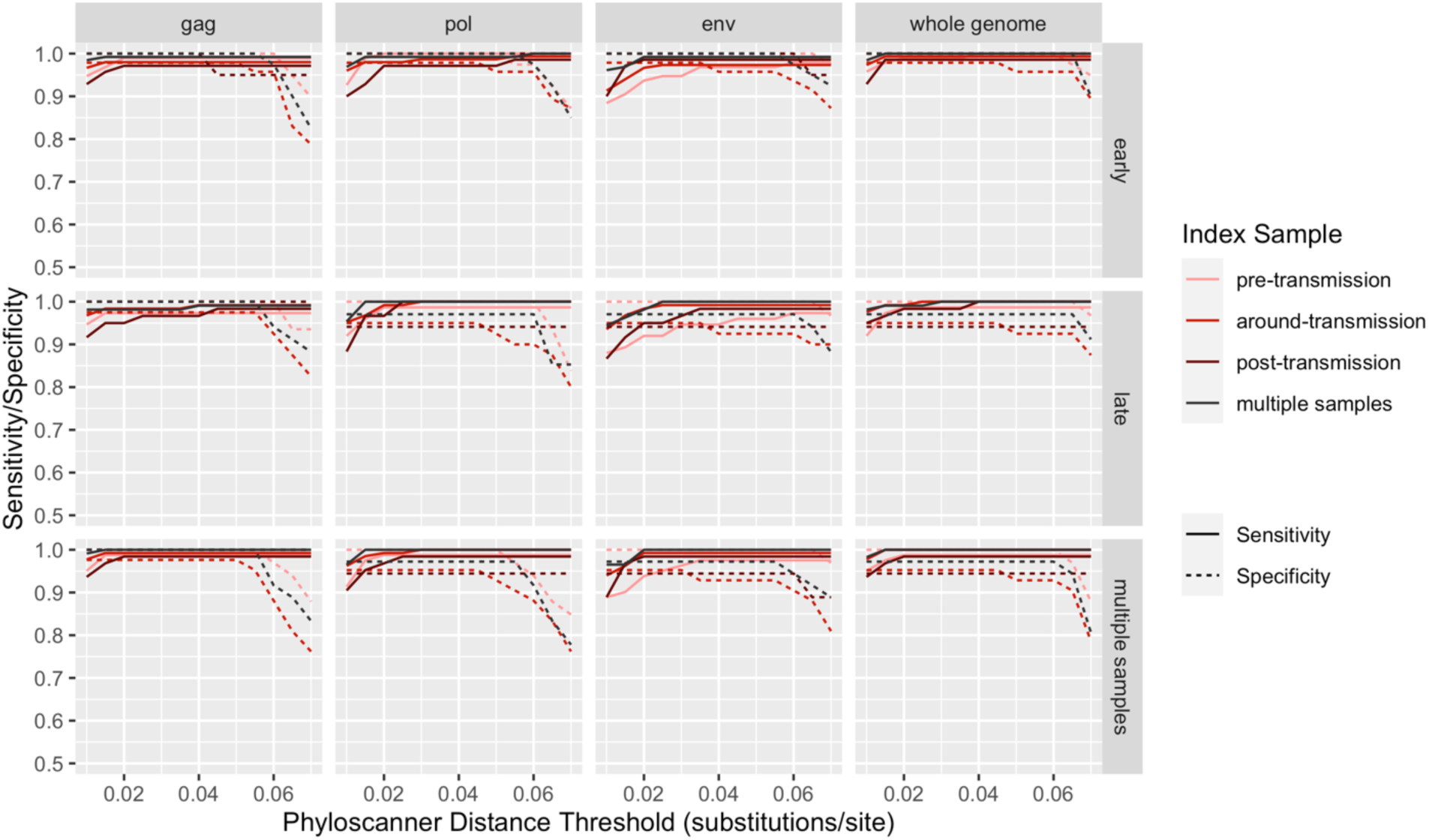
Phyloscanner sensitivity and specificity in correctly linking transmission pairs. Tests done with gag, pol, env regions and whole genome. Samples classified by sampling time with regards to the estimated time of transmission.

**Supplementary Figure S4.**
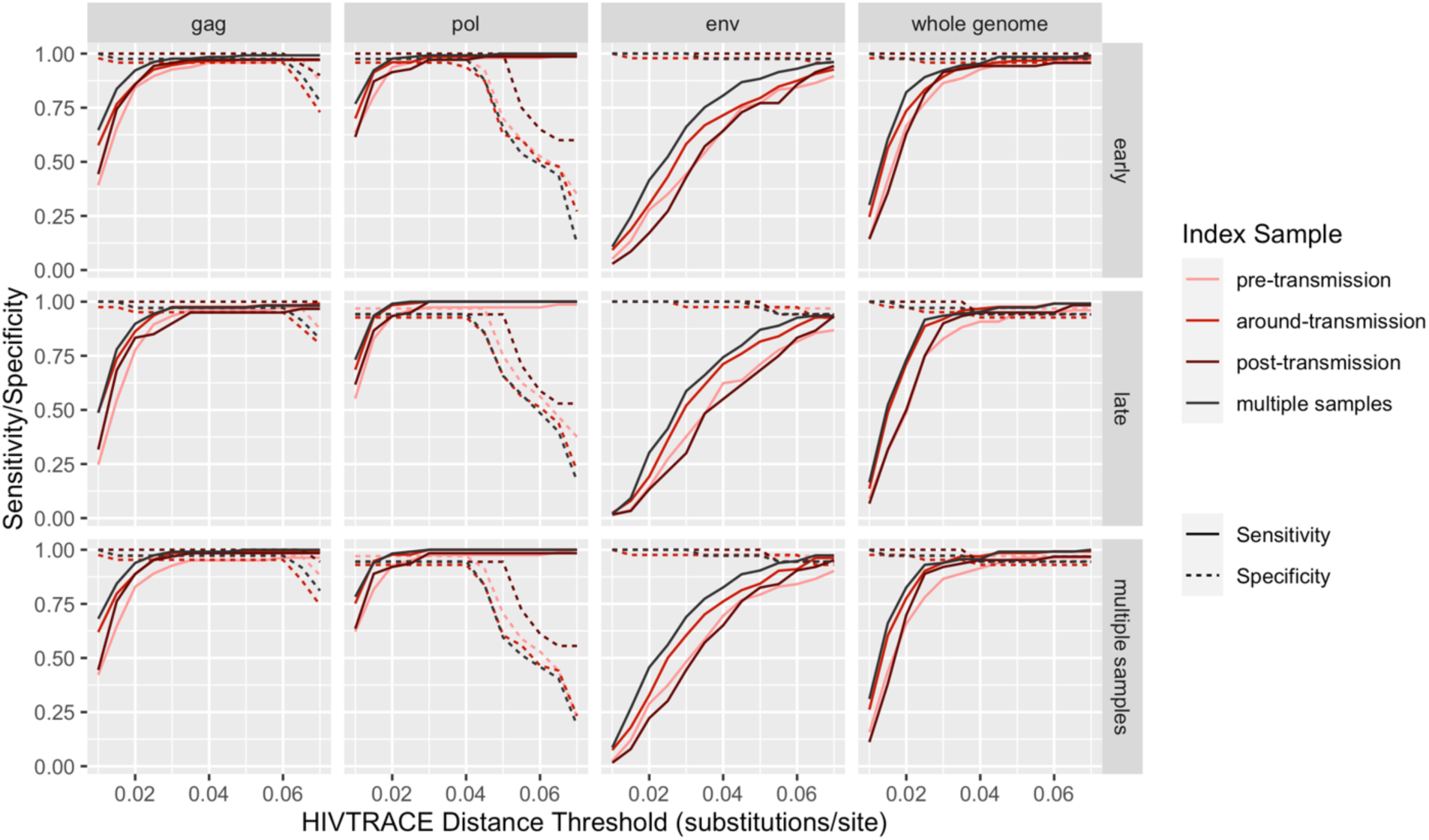
Consensus (HIV-TRACE) sequence data sensitivity and specificity in correctly clustering linked pairs. Tests done with gag, pol, env region and whole genome. Sequences classified by sampling time with regards to estimated time of transmission.

**Supplementary Figure S5.**
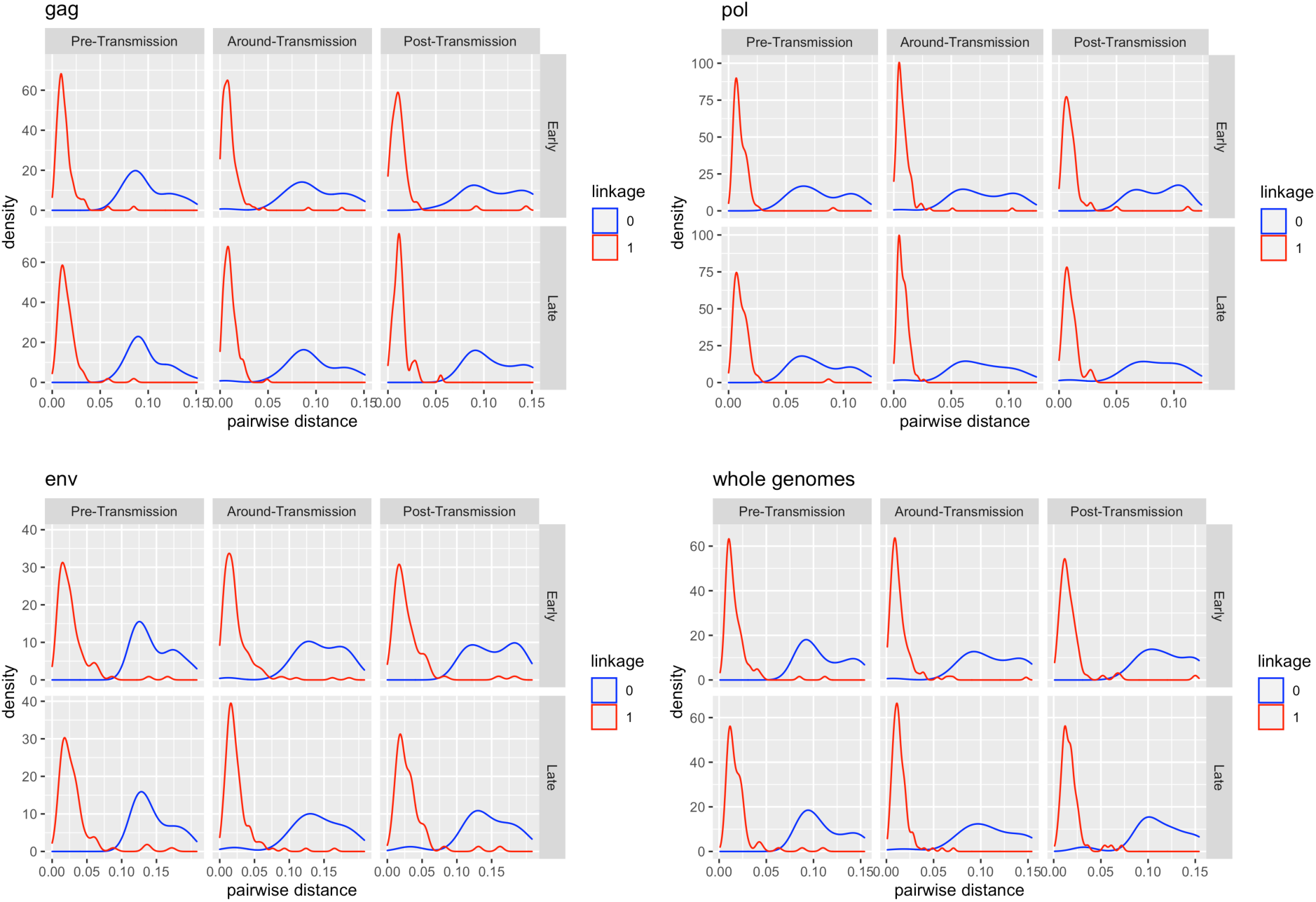
Pairwise distance distributions for all gold standard classifier assessed couples. The columns are different index sample groups (Pre-Transmission, Around-Transmission, and Post-Transmission) and the rows are different seroconverter sample groups (early and late). “1” stands for linked, “0” for not linked.

**Supplementary Figure S6.**
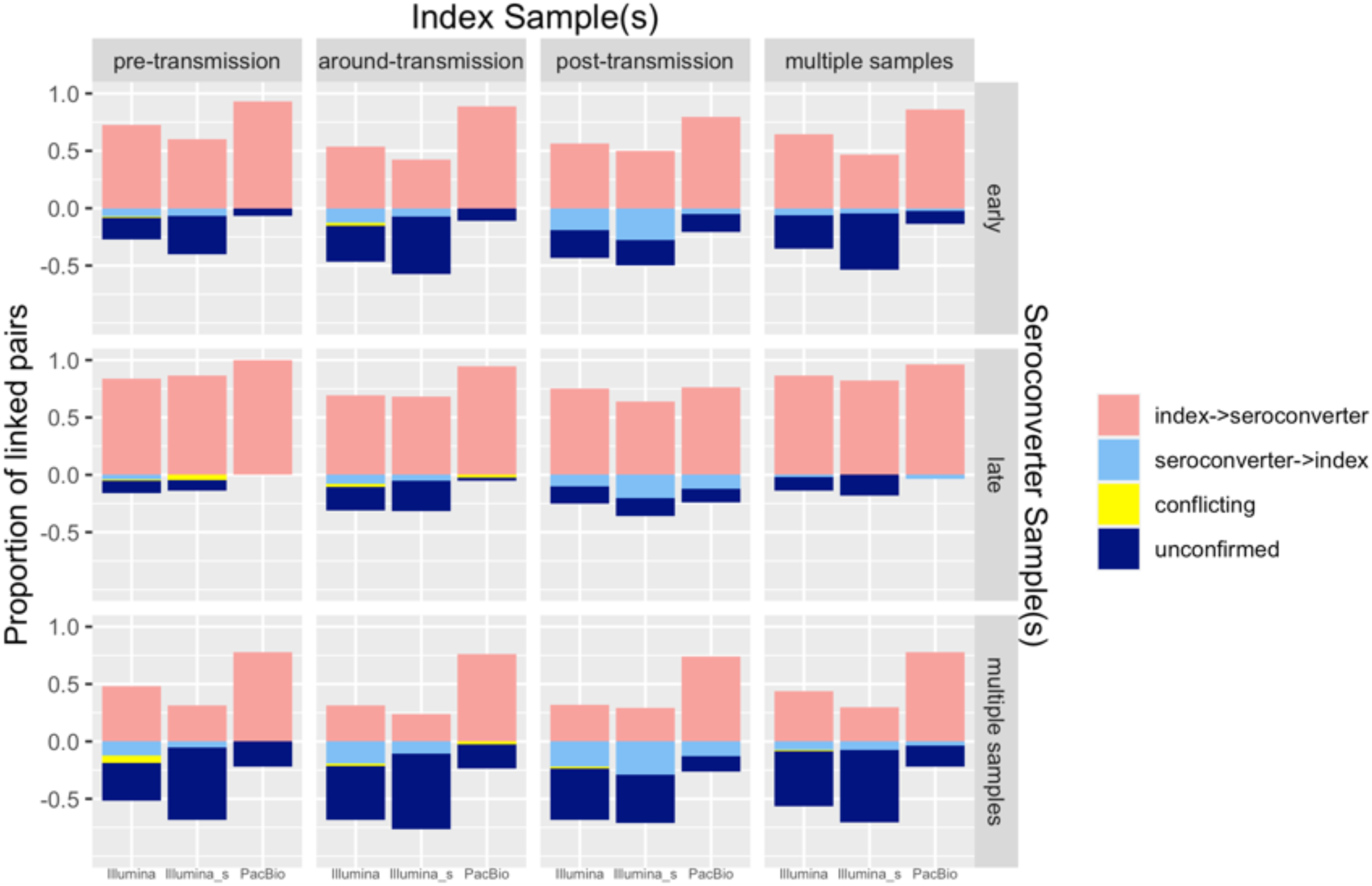
Confirmation of directionality by phyloscanner. The columns are index sampling time groups (Pre-Transmission, Around-Transmission, Post-Transmission, and multiple samples) and the rows are seroconverter sampling time groups (Early, Late, and Multiple samples). Within each box, the first column (Illumina) shows results from all available Illumina samples, the second column (Illumina-s) shows results from the subset of Illumina samples that were PacBio sequenced, the third column (PacBio) shows results from all PacBio sequenced pairs.

**Supplementary Figure S7.**
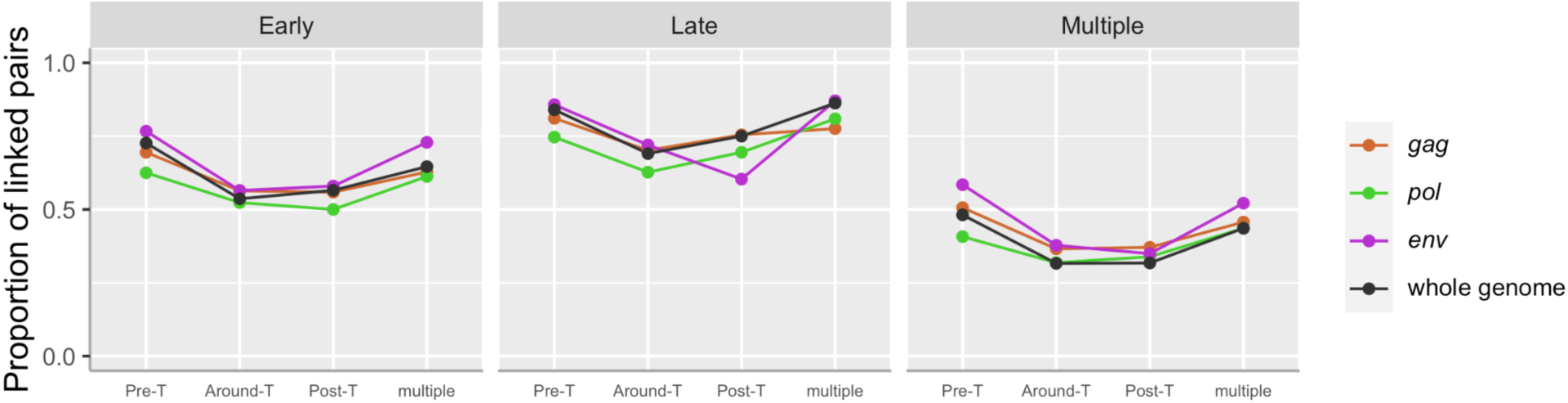
Confirmation of directionality with datasets from different genomic regions (Illumina only).

**Supplementary Figure S8.**
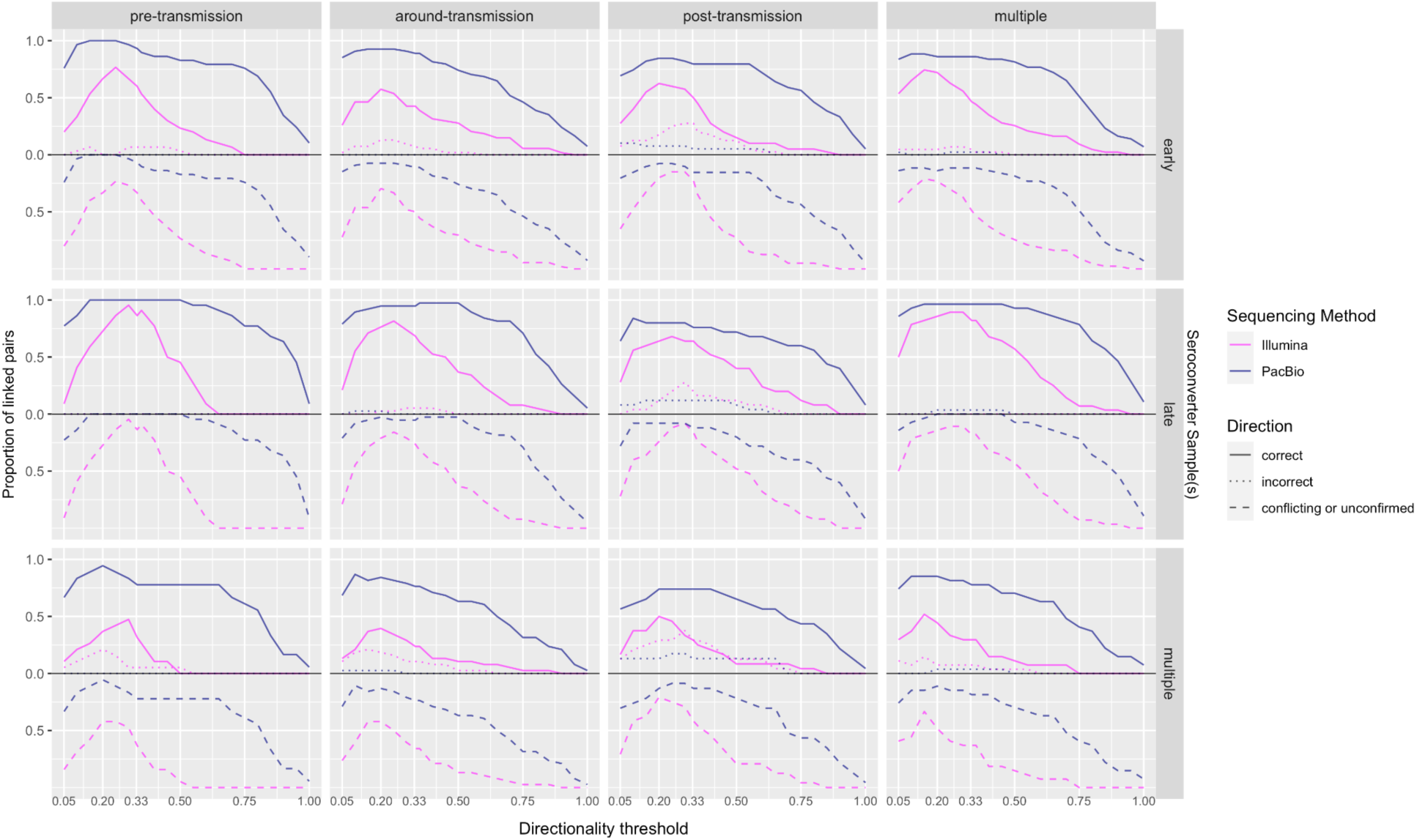
The proportion of phyloscanner linked paired with transmission direction called correctly (solid line), incorrectly (dotted line) and conflicting or unconfirmed (dashed line) across different directionality thresholds. Results from Illumina reads are coloured pink, results from PacBio reads are coloured dark blue. Proportions of correctly and incorrectly called pairs are plotted above 0, and proportion of conflicting or unconfirmed pairs are plotted below 0 to avoid overlapping lines.

**Supplementary Figure S9a&b.**
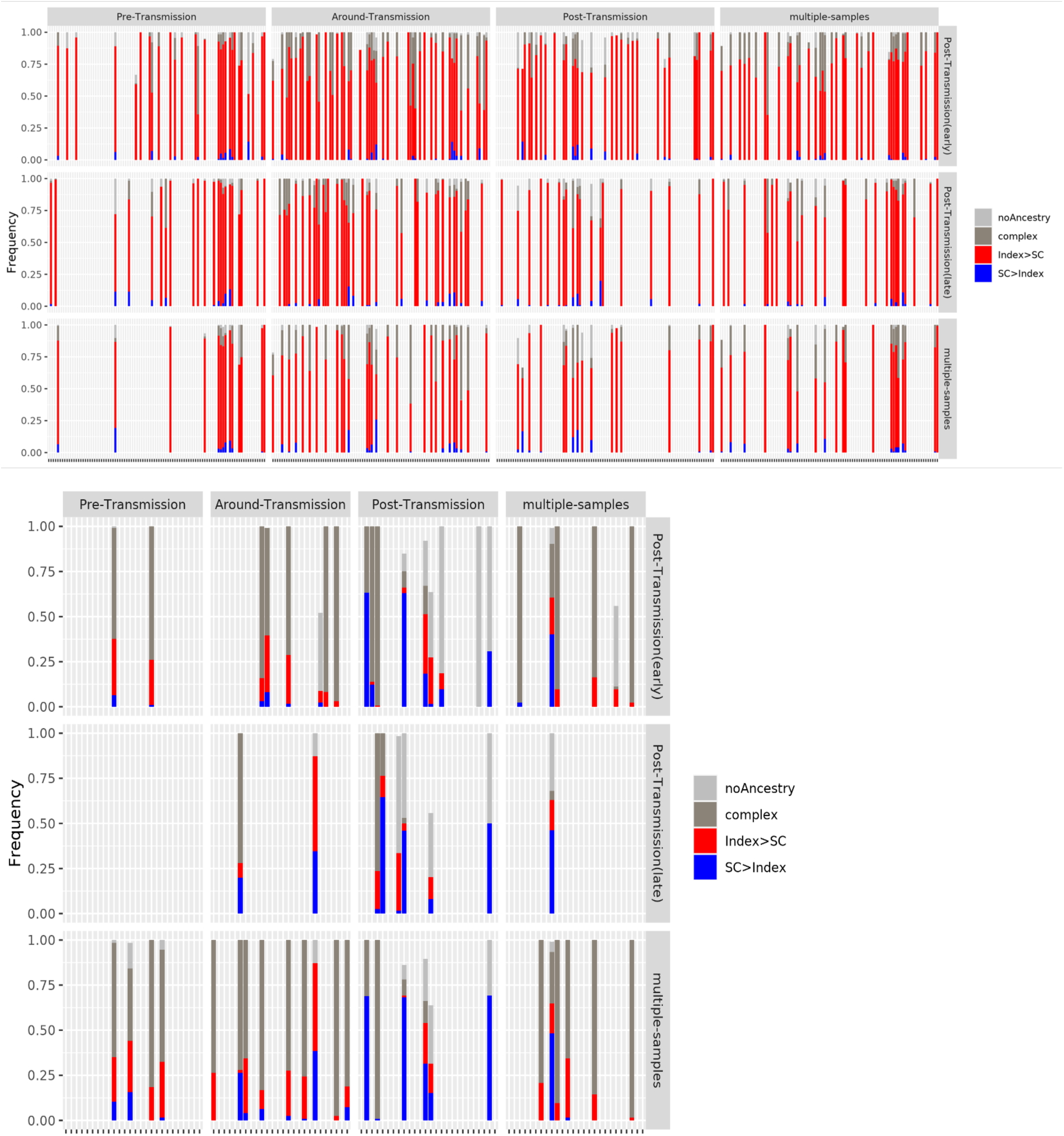
The proportion of different tree topology of all linked directed (9a) and undirected (9b) transmission pairs summarised by sample groups (PacBio only). The different tree topologies are noAncestry (light grey), complex (dark grey), index→seroconverter (red), and seroconverter→index (blue). The columns are the different index sample groups, the rows are the different seroconverter sample groups. Each tick on the x-axis is a linked transmission pair.

**Supplementary Figure S10.**
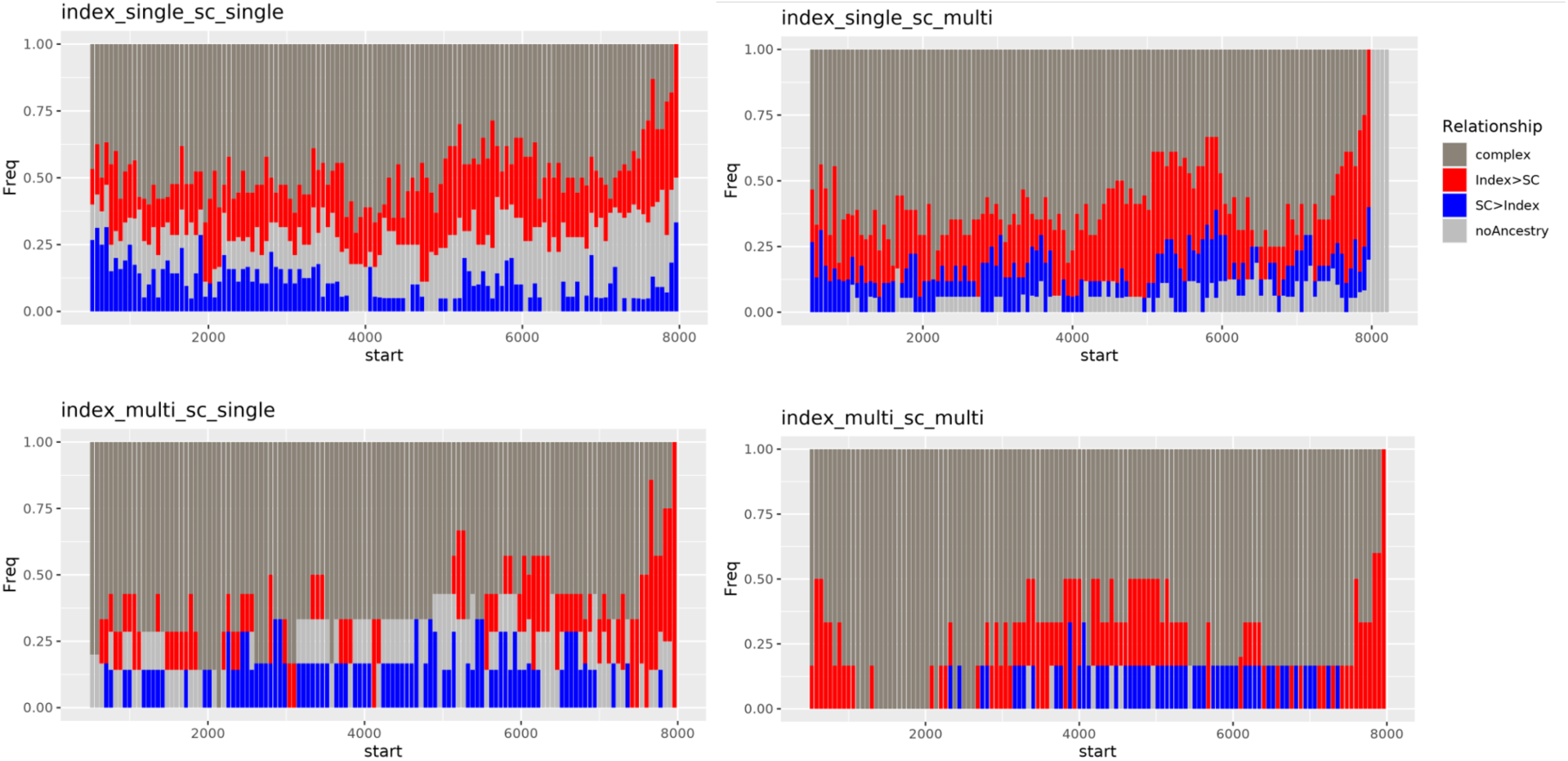
The proportion of different tree topology of all linked transmission pairs summarised across the HIV genome (PacBio only). The summaries are done for (top-left) one index-to-one seroconverter sample groups, (top-right) one index to all seroconverter sample groups, (bottom-left) all index to one seroconverter sample groups, and (bottom-right) all index to all seroconverter sample groups. The different tree topologies are noAncestry (light grey), complex (dark grey), index→seroconverter (red), and seroconverter→index (blue).

## Supplement TextFile 1

### Estimating time since infection for seroconverters in heterosexual couples

We performed one phyloscanner run for each heterosexual couple with all index and seroconverter samples sequenced with Illumina platform that generated reads covering 7500bp of the HIV-1 genome using the Phyloscanner_make_trees.py command specified below. Then we used the Phyloscanner_analyse_tree.R command specified below to produce *phyloscanner* phylogenetic summary statistics files and minor allele frequency files for all seroconverter samples. These two types of files are then used as input to HIV-phyloTSI to produce estimates of the specific sample’s time since infection.

**Phyloscanner_analyse_tree.R command** to generate phylogenetic summary statistics files for TSI estimates:

s,15 -og A1.UGANDA.2007.p191845.JX236671 -amt -sat 0.33 -sks -ow -rda -tn -v 1 -swt 0.5 - rcm -blr -pbk 15 -rtt 0.005 -rwt 3 -m 1E-5 An estimated time of transmission for each sample was calculated by subtracting the HIV-phyloTSI point estimate of time since infection from the date that sample was obtained. These were combined with the known epidemiological information (the last negative date and first positive date of the seroconverter) to estimate one time of transmission for each index-seroconverter (i.e. source-recipient) pair as follows:

a. if exactly one seroconverter sample had an estimate time between the last negative and first positive dates of that individual, that estimate;
b. or if more than one seroconverter sample had an estimate between the last negative and first positive dates, the value from the sample with the smallest estimated time since infection (where HIV-phyloTSI has less uncertainty (Golubchik et al. 2022));
c. or if all estimates fell outside the period between those two dates, the midpoint between the dates
d. or if the last negative date of the seroconverter was unavailable, their first positive date.

### Phyloscanner commands and applicable options

**Phyloscanner_make_trees.py command** for generating sliding window alignments:

-P -A reference_alignment.fasta -2 HXB2_K03455 -XR HXB2_K03455 -XC [DRsites] --min-read-count 1 --x-mafft mafft --no-trees

Masked drug resistance sites:

[DRsites]:

823,824,825,892,893,894,907,908,909,1012,1013,1014,1156,1157,1158,1384,1385,1386,1444, 1445,1446,1930,1931,1932,1957,1958,1959,2014,2015,2016,2023,2024,2025,2080,2081,2082, 2134,2135,2136,2191,2192,2193,2280,2281,2282,2283,2284,2285,2298,2299,2300,2310,2311, 2312,2316,2317,2318,2319,2320,2321,2322,2323,2324,2340,2341,2342,2346,2347,2348,2349, 2350,2351,2352,2353,2354,2355,2356,2357,2358,2359,2360,2373,2374,2375,2379,2380,2381, 2385,2386,2387,2388,2389,2390,2391,2392,2393,2394,2395,2396,2400,2401,2402,2409,2410, 2411,2412,2413,2414,2415,2416,2417,2424,2425,2426,2430,2431,2432,2436,2437,2438,2439, 2440,2441,2442,2443,2444,2457,2458,2459,2460,2461,2462,2463,2464,2465,2469,2470,2471, 2472,2473,2474,2478,2479,2480,2481,2482,2483,2496,2497,2498,2499,2500,2501,2502,2503, 2504,2505,2506,2507,2514,2515,2516,2517,2518,2519,2520,2521,2522,2526,2527,2528,2529, 2530,2531,2535,2536,2537,2670,2671,2672,2679,2680,2681,2703,2704,2705,2709,2710,2711, 2733,2734,2735,2742,2743,2744,2748,2749,2750,2751,2752,2753,2754,2755,2756,2757,2758, 2759,2769,2770,2771,2772,2773,2774,2778,2779,2780,2811,2812,2813,2814,2815,2816,2817, 2818,2819,2823,2824,2825,2841,2842,2843,2847,2848,2849,2850,2851,2852,2856,2857,2858, 2865,2866,2867,2871,2872,2873,2892,2893,2894,2895,2896,2897,2901,2902,2903,2904,2905, 2906,2952,2953,2954,2961,2962,2963,3000,3001,3002,3015,3016,3017,3018,3019,3020,3030, 3031,3032,3042,3043,3044,3084,3085,3086,3090,3091,3092,3099,3100,3101,3111,3112,3113, 3117,3118,3119,3135,3136,3137,3171,3172,3173,3177,3178,3179,3180,3181,3182,3189,3190, 3191,3192,3193,3194,3204,3205,3206,3210,3211,3212,3222,3223,3224,3228,3229,3230,3237, 3238,3239,3246,3247,3248,3249,3250,3251,3255,3256,3257,3261,3262,3263,3396,3397,3398, 3501,3502,3503,3546,3547,3548,3705,3706,3707,4425,4426,4427,4449,4450,4451,4503,4504, 4505,4518,4519,4520,4590,4591,4592,4641,4642,4643,4647,4648,4649,4656,4657,4658,4668, 4669,4670,4671,4672,4673,4692,4693,4694,4722,4723,4724,4782,4783,4784,4974,4975,4976, 5016,5017,5018,5067,5068,5069,7863,7864,7865,7866,7867,7868,7869,7870,7871,7872,7873, 7874,7875,7876,7877,7881,7882,7883,7884,7885,7886

### Phyloscanner_analyse_tree.R command to for transmission pair analysis

s,15 -og A1.UGANDA.2007.p191845.JX236671 -m 1E-5 --RDAonly -ow -ns -pbk 15 -rwt 3 -rtt 0.01 -rcm -swt 0.5 -sdt 0.02 -amt -sat 0.33 -v 1

13 Phyloscanner alignment **reference sequence GenBank ID**s:

Subtype A1 (JX236671;KU168256;KY658714;AB253429;KT152842);

subtype C (AF443088;AY228557;AY162223;KC156218);

subtype D (JX236672;KU168271;AJ320484);

subtype B(K03455).

### HIV-TRACE options

In addition to the options specified in the manuscript, **other options used for all HIV-TRACE**

runs: -a resolve -m 500 -g .05

